# A New Pathway for the Decussation of Corticospinal Tracts

**DOI:** 10.1101/2025.06.03.657394

**Authors:** Hsiu-Ling Li, Mohamed Ahmed, Nahla Zaghloul

## Abstract

Each brain hemisphere controls the movements of the opposite side of the body because the motor axons cross the midline, known as CST decussation. The current theory on CST decussation suggests CSTs decussate as a single tract at the junction between medulla and spinal cord. Although this theory is widely accepted, this theory is based on selective analyses and is therefore incomplete. Here, we employed new approaches, including the horizontal analyses and a non-invasive anterograde tracing method to examine CST decussation thoroughly. We analyzed all CS axons in 3 planes. These approaches led to the discovery of a new pathway for CST decussation. We found CSTs turned back to medulla, moving anteriorly and decussated in an oval structure. In this structure, each CST split into 4 fascicle groups and interdigitated with the corresponding groups from the opposite CST to cross the midline. The significance of this pathway was apparent after decussation where these 4 groups reversed direction, moving posteriorly toward spinal cord. While moving, the motor axons gradually separated at different locations and subsequently turned and occupied the correct positions in the dorsal funiculus for proper limb control. In addition to CSTs, we also characterized several components in the oval structure.

## Introduction

Corticospinal tracts (CSTs) originating from the primary motor cortex (M1) and terminating in the spinal cord, control the voluntary movements of the limbs/trunk. However, only a small number of CST fibers stay on the same side. The majority of CST fibers cross the midline at the medulla and enter the opposite side of the spinal cord. CST decussation has been extensively studied ^1,2^, but none of these reports provided a complete understanding of CST decussation because they did not perform an analysis in the horizontal plane and they only studied selective CST fibers.

In contrast to previous reports, we examined CST decussation in the horizontal plane and included all CS fibers in our analysis using 1) PKC-γ immunostaining and 2) a novel tracing method that traced the neuraxis *in vitro*. These approaches, unexpectedly, revealed a new pathway for CST decussation which was subsequently verified in the coronal and sagittal planes. We found CSTs did not decussate at the junction between medulla and spinal cord. Instead, CSTs turned over 90^0^ to face the medulla again and then moved anteriorly above the pyramidal tracts to enter an oval structure surrounding the midline. Decussation occurred in this structure. During decussation, each CST split into 4 fascicle groups. Each group then interdigitated with the corresponding groups from the opposite CST to cross the midline. The significance of this pathway was apparent after decussation. After crossing the midline, the 4 fascicle groups changed direction, moving posteriorly toward spinal cord. While moving, the CS fibers in each fascicle group separated at different locations. This separation allowed each CS fiber to turn and occupy the correct positions in the dorsal funiculus and that is critical for proper limb control. Please note, this turning process was seen in the coronal and sagittal sections, not in the horizontal sections.

Coronally, the oval structure was sectioned along its length. Thus, CST decussation and separation could be seen in a series of coronal sections that spanned over 500 μm. As stated above, the separated CS axons were seen to turn and enter the dorsal funiculus at different locations. Sagittally, the oval structure appeared as a belt-like structure. CST could be seen to split/decussate in this structure and separate after decussation. The separated CS fibers subsequently followed a turning strategy to enter the dorsal funiculus in an orderly manner. We found the CS fibers that separated anteriorly (A/P coordinates), turned laterally (M/L coordinates) and entered the dorsal positions (D/V coordinates) of the dorsal funiculus and vice versa.

Together, this new pathway showed the utility of the oval structure in CST decussation, which not only assisted CS axons to cross the midline, but also allowed each CS axon to separate at different locations in order to enter the correct positions in the dorsal funiculus.

Apart from CSTs, we also characterized the oval structure. We found GFAP^+^ cells led CSTs to enter the oval structure. The smooth muscle actin^+^ cells were the main residential cells inside the oval structure, whereas the non-CST nerve fibers were the main components in the concentric layers surrounding the oval structure.

## Results

In this paper, this new pathway for CST decussation is presented from the views of 3 anatomical planes. The CSTs in each plane were analyzed by two methods: 1) PKC-γ immunostaining and 2) a novel anterograde tracing. To assist in reading these analyses, each dataset is preceded by a diagram that outlines the experimental design and the appearance of CSTs.

### 1. Horizontal Analysis

#### Method (I): PKC Immunostaining

For PKC-γ immunostaining, we used a tissue section that contained the entire medulla and 3-5 segments of spinal cord (top diagram, Fig. 1A). This tissue was sectioned in each of the 3 planes and then immunostained for PKC-γ. Specific staining on CSTs was seen in the horizontal sections, but not in the coronal or sagittal sections.

**Figure 1:**
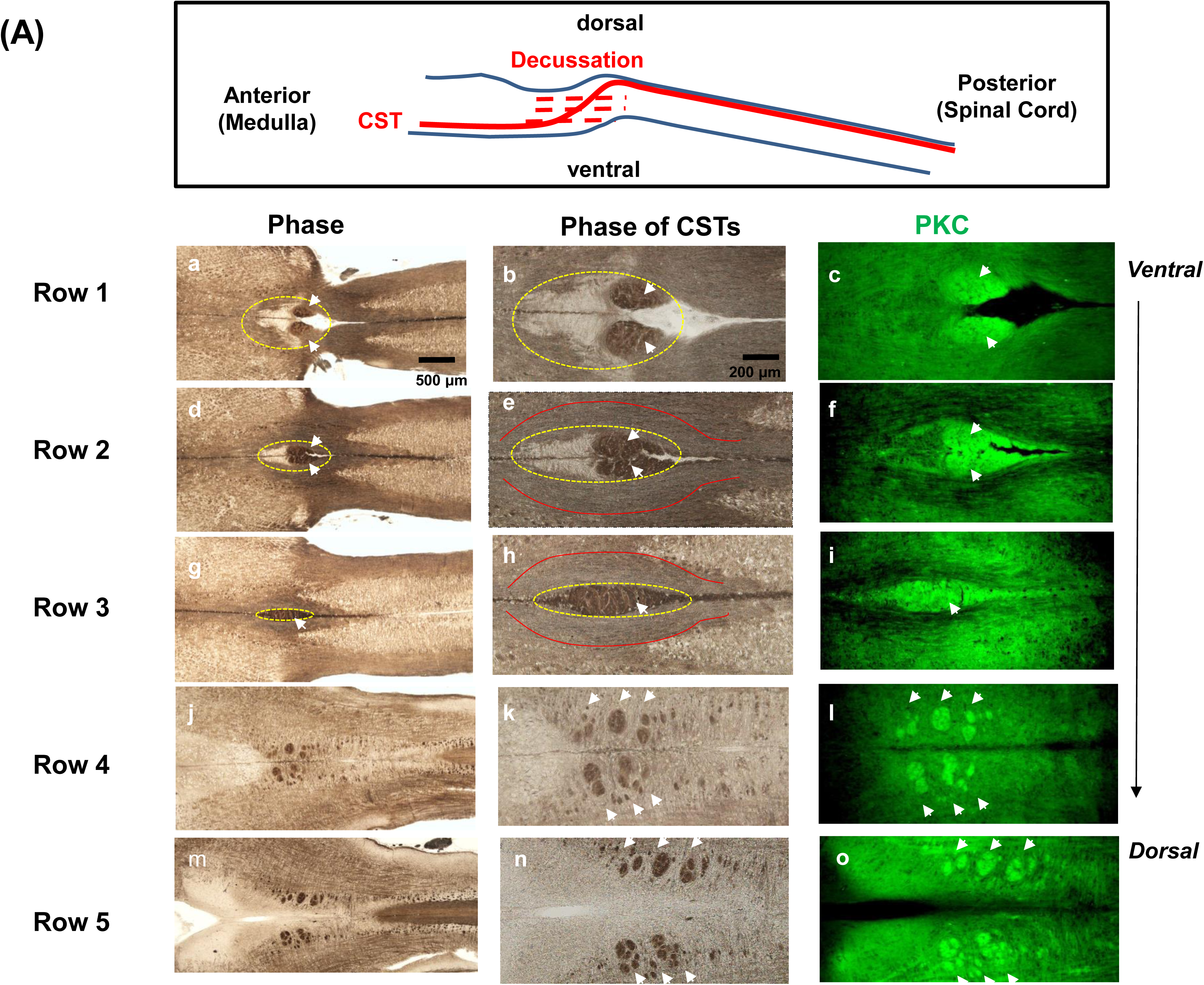
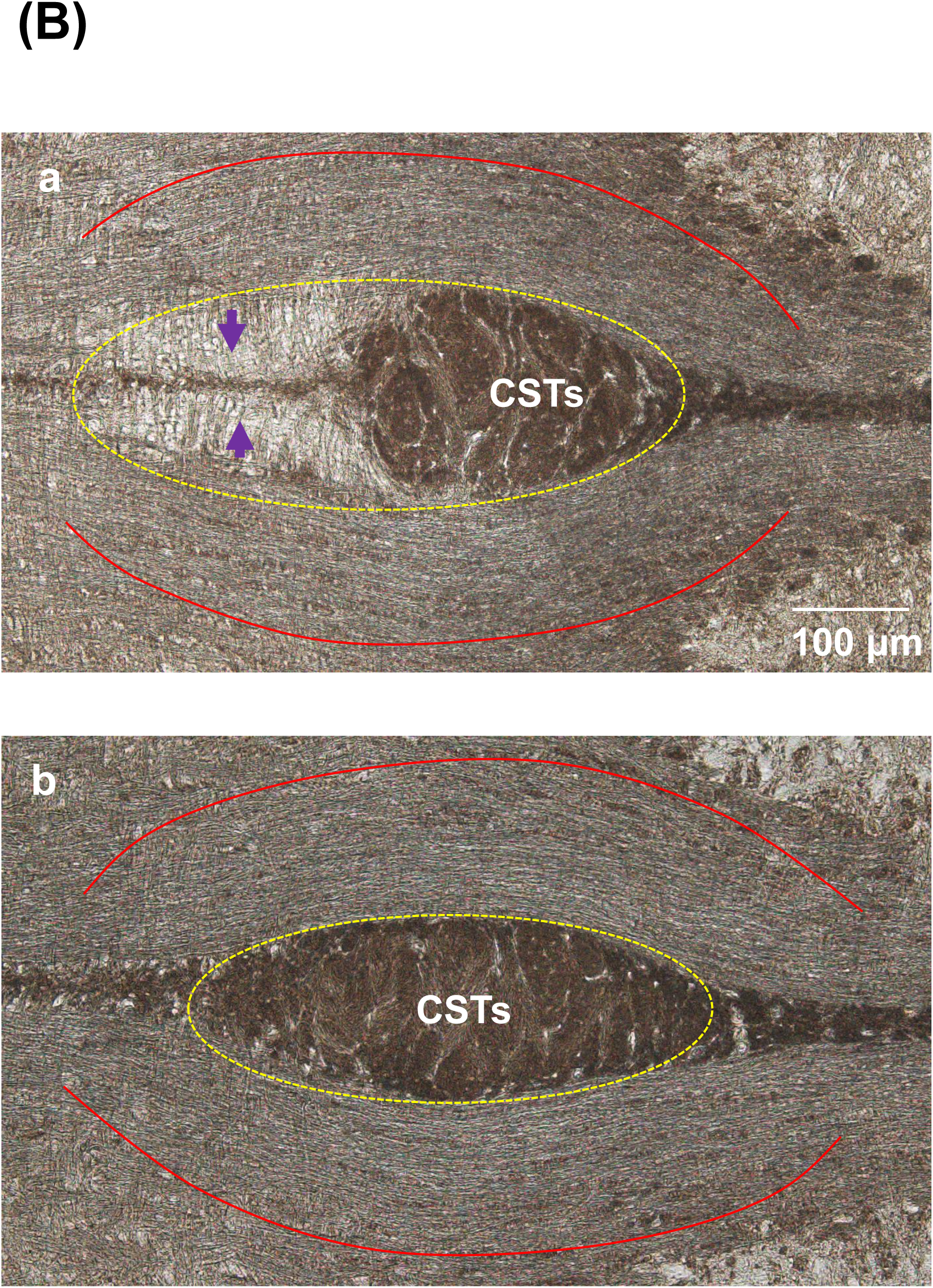
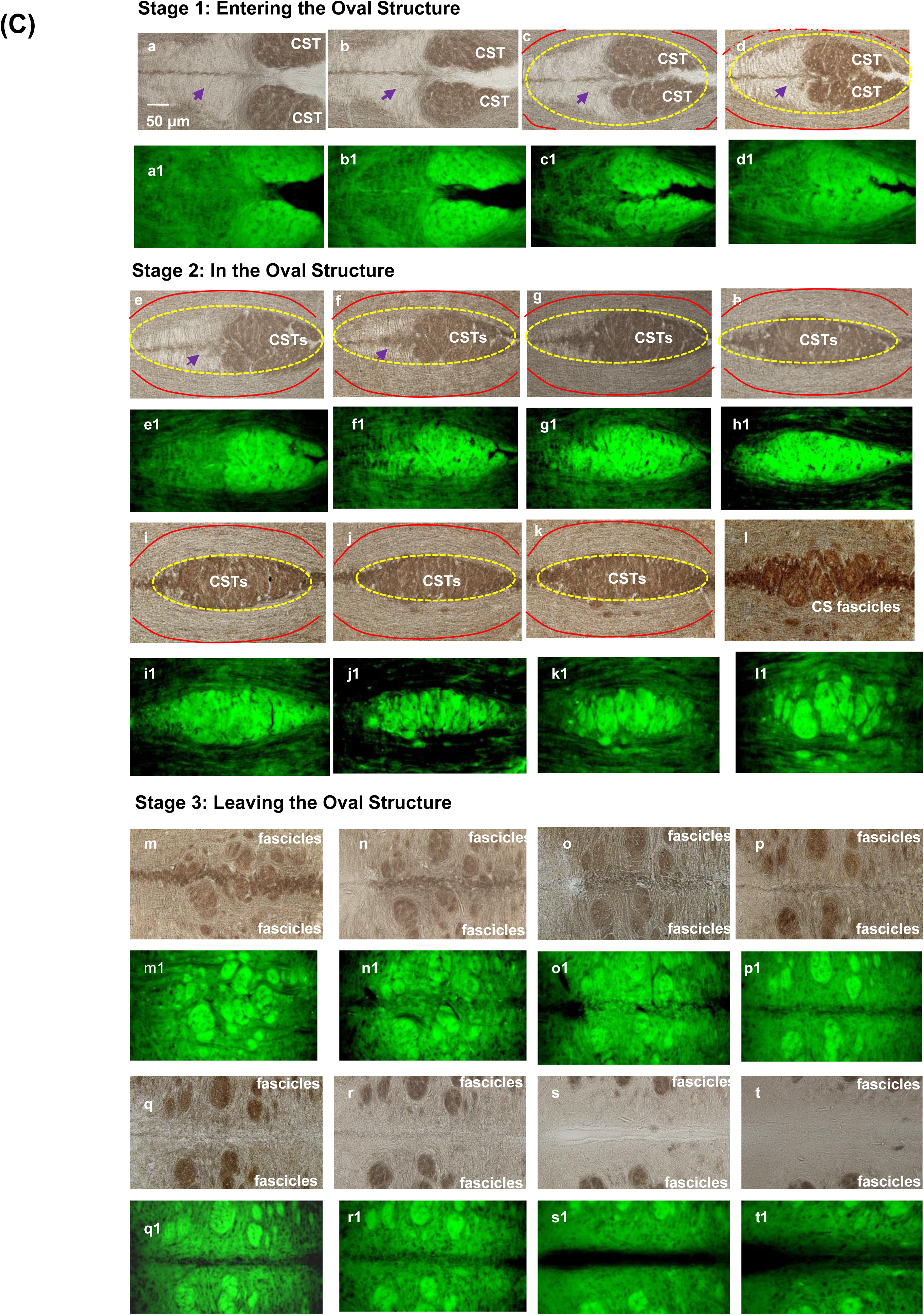
Horizontal analysis of CST decussation by PKC-γ immunostaining. **(A)** CST decussation is summarized under low magnification, which shows the experimental design (top) and the data from 5 horizontal sections (bottom) (n>100). Each row is one section. Each section has 3 images; the phase of the entire section (left), the phase of the area surrounding CSTs (middle) and the results of PKC-γ immunostaining (right). CSTs (white arrows) move back to medulla, enter an oval structure (dashed yellow lines, a-f) and become a long tract at midline (g-i). The oval structure is sandwiched by numerous concentric lines on both sides (outlined by 2 red curves). CSTs leave the structure as multiple fascicles (j-o). **(B)** Enlarged phases which show the details of the oval structure (dashed yellow lines), including the optically dense lines (purple arrows) and the concentric layers externally (outlined by 2 red curves) **(C)** CST decussation in the oval structure is detailed under high magnification. Each horizontal section is represented by a pair of images; the phase on the top (e.g. a) and the result of PKC-γ immunostaining at the bottom (e.g. (a1). The oval structure and the residential cells inside are shown by dashed yellow lines and purple arrows (a-f). The 2 red curves outline the concentric layers surrounding the oval structure (a-k).

Figure 1A summarizes the horizontal analysis of CST decussation in 5 horizontal sections from ventral to dorsal sections under low magnification. Each row is one section and each row has 3 image columns which show the phase of the entire section (left), the phase of the area surrounding the CSTs and the result of PKC-γ immunostaining (middle and right columns). Figure 1A begins with a very ventral section in row 1. The phase of this section is shown in (a), in which the wider area is medulla and the narrow area is spinal cord.

At their junction, there is a pair of optically dense spheres (white arrows), facing an oval structure (dashed yellow lines). PKC-γ immunostaining suggested that this pair of spheres were CSTs. The phase of the area around the CSTs and the corresponding result of PKC-γ immunostaining are shown in (b) and (c). We then move up to the next dorsal level in row 2. However, there is no sign of CST decussation. Instead, CSTs move back to medulla, traveling antero-dorso-medially to enter the oval structure (dashed yellow lines). In row 3, the two CSTs fully occupy this structure and merge into one long tract at midline, suggesting CST decussation may occur inside this oval structure. Externally, the oval structure is enclosed by numerous concentric lines on both sides (outlined by 2 red curves). In a more dorsal section in row 4, CSTs leave the oval structure, separating into more fascicle groups and reversing the direction to travel latero-dorso-posteriorly toward spinal cord (white arrows in rows 4 and 5). The fate of these separated fascicles could not be determined in horizontal sections since they were found to turn and enter the dorsal funiculus in coronal and sagittal sections.

Given the importance of the oval structure, the details of this structure and CSTs are shown in the enlarged phase images in Figure 1B. Prior to decussation, CSTs move anteriorly to enter the oval structure (a), which is occupied by optically dense lines (purple arrows) and whose size is about 450 μm in length and 200 μm in width. However, when CSTs fully occupy this structure (b), the optically dense lines disappear and this structure becomes longer and narrower (about 500 μm in length and 180 μm in width). Externally, the oval structure is surrounded by numerous concentric layers (represented by 2 red curves in (a) and (b)).

Figure 1C shows CST decussation in the oval structure under high magnification. In Figure 1C, each horizontal section is represented by a pair of images. The phase of each section is at the top and the corresponding result of PKC-γ immunostaining is at the bottom. Consistently, before CSTs enter the oval structure (stage 1), this structure is wider and the interior is divided by optically dense lines (dashed yellow lines and purple arrows). However, when CSTs enter the oval structure (stage 2), this structure becomes narrower and the space with optically dense lines retracts and finally disappears. Based on the changes in size, this oval structure looks like an oval-shaped pyramid. In contrast to the interior, the oval structure is externally surrounded by numerous concentric lines on both sides (represented by 2 red curves). When CSTs leave the oval structure after decussation (stage 3), the oval structure gradually disappears and CSTs become several fascicle groups, moving latero-dorso-posteriorly toward spinal cord.

#### Method (II): Non-invasive Anterograde Tracing

We next investigated CST decussation by tracing. However, none of the current tracing methods could label all CST fibers easily due to the invasive procedures and the variability in labeling results ^3^. We therefore developed a non-invasive tracing method to label all CST fibers. Briefly, a DiI solution was prepared and backfilled into a filamented borosilicate glass capillary and injected into one side of pyramidal tracts or M1 of a fixed mouse neuraxis *in vitro* using a stereotaxic apparatus (Figure 2A). The injected DiI labeled one CST by anterograde diffusion and the signals were stronger when injected into one pyramidal tract. Using this method, both retrograde and trans-synaptic diffusion of DiI were negligible. The labeling specificity of this method was evaluated in the horizontal analysis. Moreover, DiI was used for this tracing method, not DiO and DiD due to their limited diffusion.

**Figure 2:**
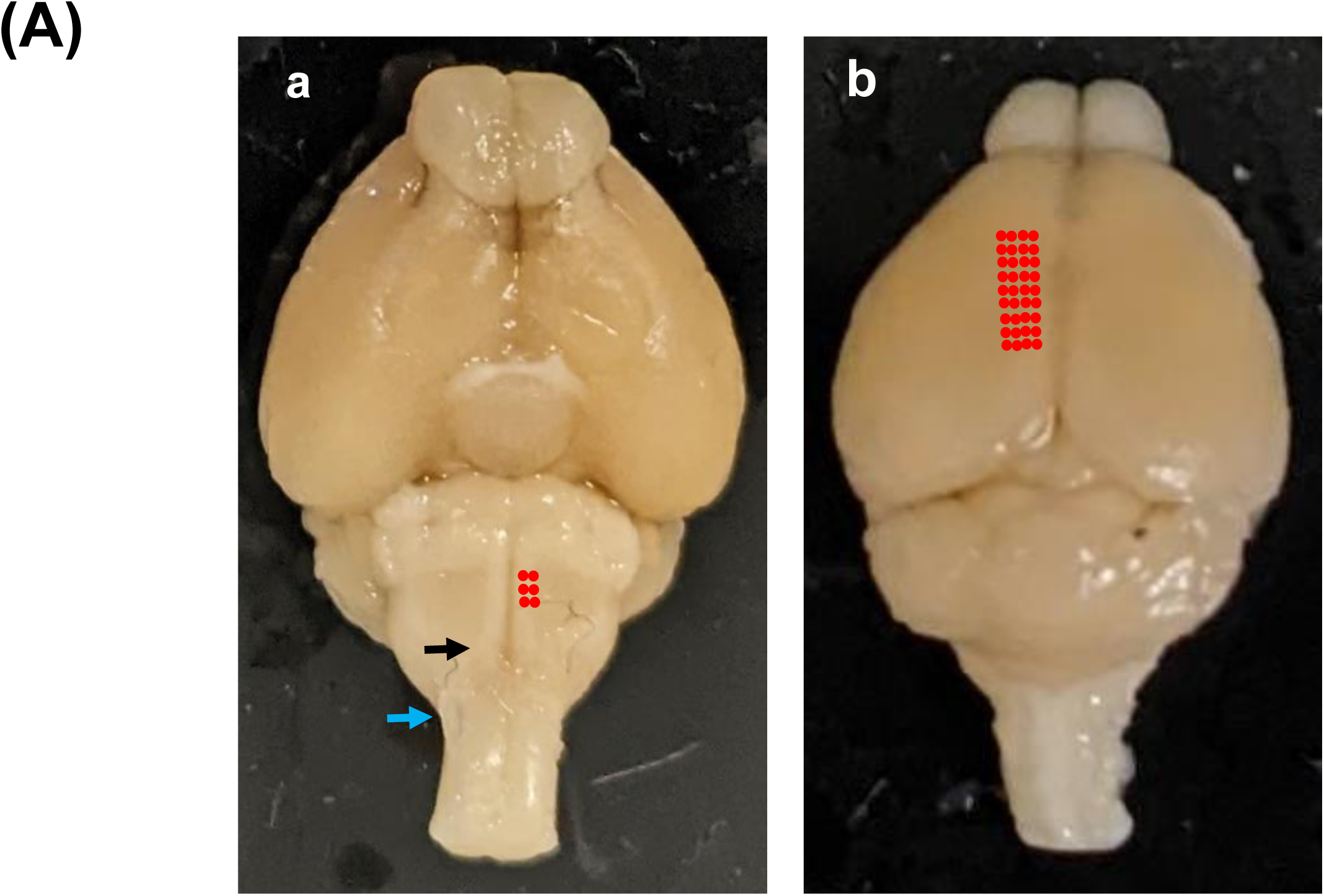
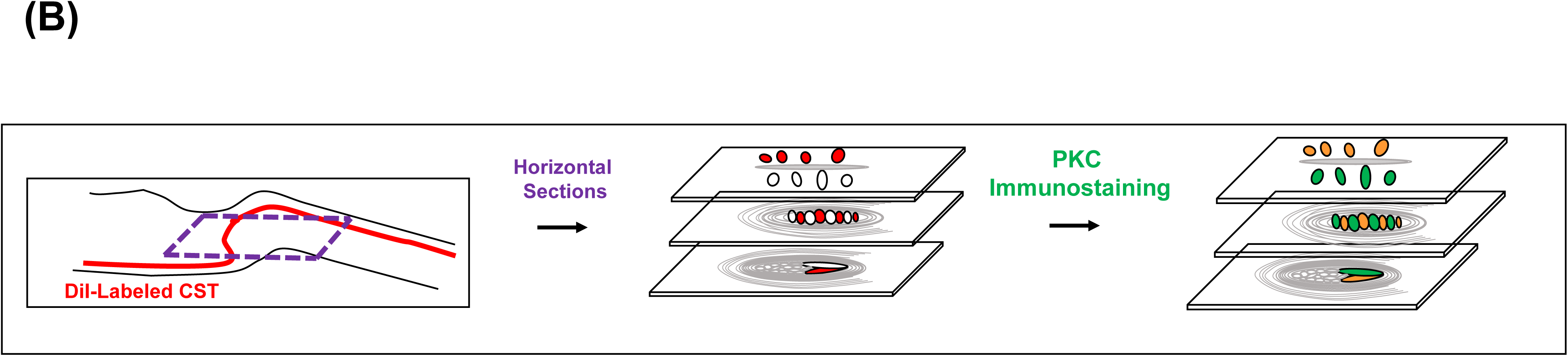
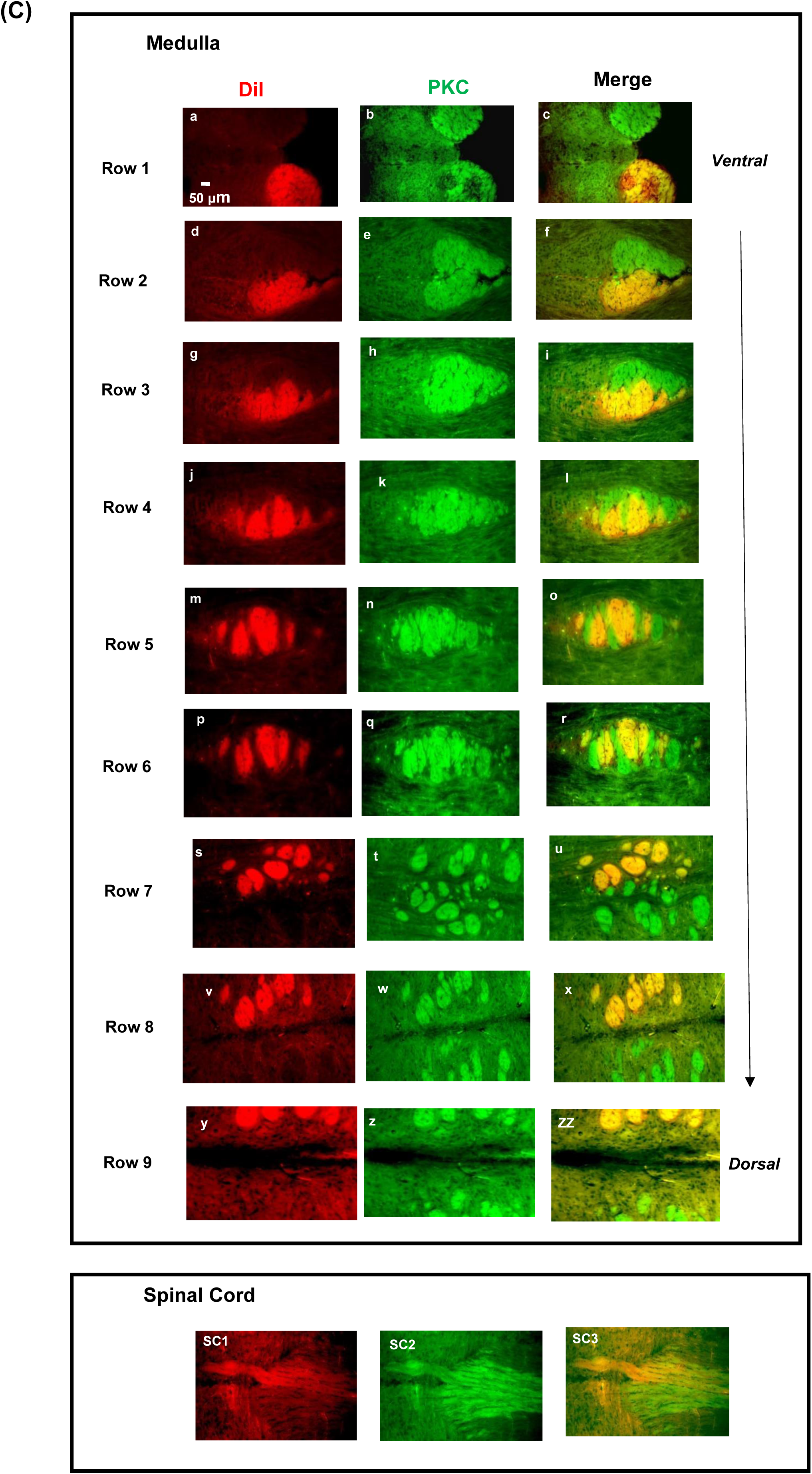
Horizontal analysis of CST decussation by DiI tracing. (A) The DiI-injection sites in one pyramidal tract and unilateral M1 (red, a, b), the junction between medulla and spinal cord (blue, b) and the location of the decussation (black, b). (B) This diagram outlines the horizontal analysis. (C) The results of 10 horizontal sections where each row is one section in ventral to dorsal order (n>10). Each section has 3 images; the results of DiI (red, left column), PKC-γ immunostaining (green, middle column) and merged (right column). CSTs move back to medulla to enter an oval structure (c,f, rows 1-2). Each CST splits into 4 fascicle groups and interdigitates with their counter groups from the other CST to cross the midline (g-r, rows 3-6). After decussation, CSTs leave the oval structure as multiple fascicles and move toward spinal cord (u,x,zz, rows 7-9). CSTs moves horizontally from the dorsal funiculus of medulla to that of spinal cord (SC1-3).

In the horizontal analysis, the DiI-labeled neuraxis was sectioned horizontally and then immunostained for PKC-γ (diagram in Fig. 2B). This approach allowed us to evaluate this tracing method and examine CST decussation simultaneously. Figure 2C shows the results of 9 horizontal sections from ventral to dorsal sections. In this figure, each row is one horizontal section and each section has 3 image columns. To evaluate this tracing method, we compare these 3 columns. The DiI-labeled CST is shown in red (left column) whereas PKC-γ immunostained both CSTs in green (middle column). When merged (right column), the DiI-labeled tract is also positive for PKC-γ immunostaining and therefore appears orange/yellow. In contrast, the unlabeled tract only showed the green color of PKC-γ immunostaining. These results validated the specificity of our tracing method.

To analyze CST decussation, we compare the results from different rows, i.e. from ventral to dorsal sections. In rows 1 and 2, CSTs move back to medulla, travelling antero-dorso-medially to enter an oval structure. As the two tracts move closer to the midline in the oval structure, each CST gradually splits into 4 fascicle groups (rows 3 and 4). Each group then interdigitates with the corresponding group from the other CST to cross the midline (rows 5 and 6). This process is associated with increased extracellular space as shown by the increased expression of hyaluronic acid (supplementary Fig. S1). After decussation, these 4 groups change direction, moving latero-dorso- posteriorly, toward the spinal cord (rows 7, 8 and 9). While moving, the CS axons in these 4 groups gradually separate. The fate of these separated CS axons cannot be seen on horizontal sections since these axons are shown to turn and enter the dorsal funiculus of medulla in coronal and sagittal sections. After turning, the CS axons in the dorsal funiculus of medulla move horizontally to that of spinal cord (SC1-3, Fig. 2C) suggesting the CS axons maintained the same positions in both structures. Thus, the CS topography in the spinal cord which is critical for proper limb control results from CST decussation in medulla.

### 2. Coronal Analysis

*Method (I): PKC Immunostaining:* N/A due to the lack of specificity.

#### Method (II): Non-Invasive Anterograde Tracing

To assist in reading the coronal analysis, we drew a diagram to outline this analysis in Fig. 3A. Here, the DiI-labeled neuraxis was sectioned coronally and analyzed without PKC-γ immunostaining. Figure 3B shows the results from anterior to posterior sections. In contrast to previous reports ^1,2^, CST decussation can be seen in a series of coronal sections that span over 500 μm. However, the appearance and the intensity of the DiI signals in each section are different. In the most anterior sections (a-c), the DiI signals are faint (yellow arrows) and the decussation occurs on top of the pyramidal tracts (green arrows). Stronger DiI signals can be seen in the more posterior sections (d-n). However, when close to the completion of decussation, the DiI signals in the dorsal funiculus gradually become intense (blue arrows in o-w). Moreover, the DiI-labeled CST is seen as a single DiI tract on the ipsilateral side, but separates into several CST fascicles when crossing the midline to the contralateral side. Each of the separated fascicles is localized at different positions and therefore turns and enters the dorsal funiculus at different A/P and M/L coordinates (blue arrows). This separation also determines the D/V coordinates since the dorsal funiculus are at different heights in different sections. However, this phenomenon is much clearer on the sagittal sections shown below (Fig. 3D). Together, these coronal data not only support our new model for CST decussation, but also reveal the significance of this pathway in determining the positions of CS axons in the dorsal funiculus.

**Figure 3:**
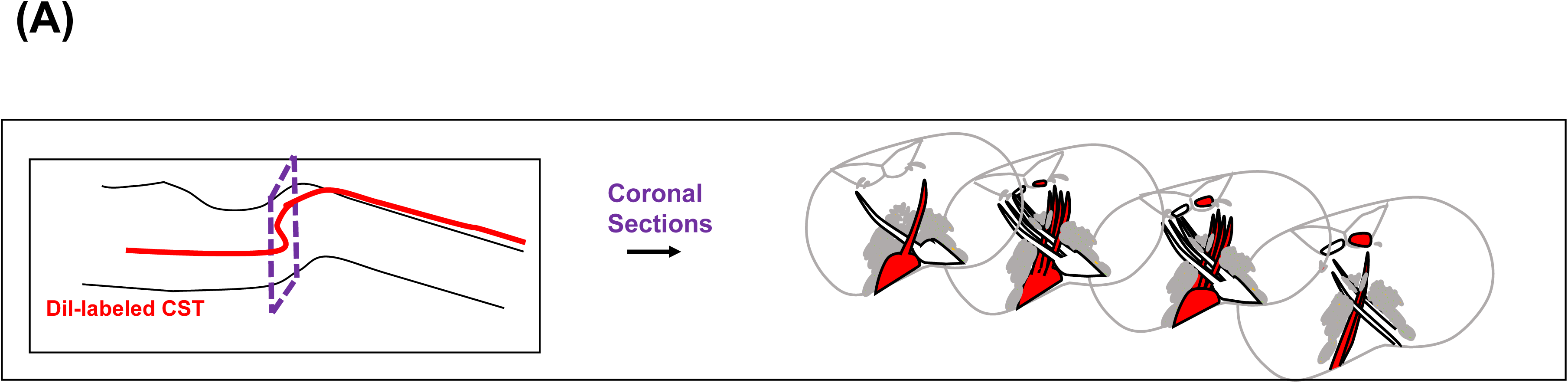
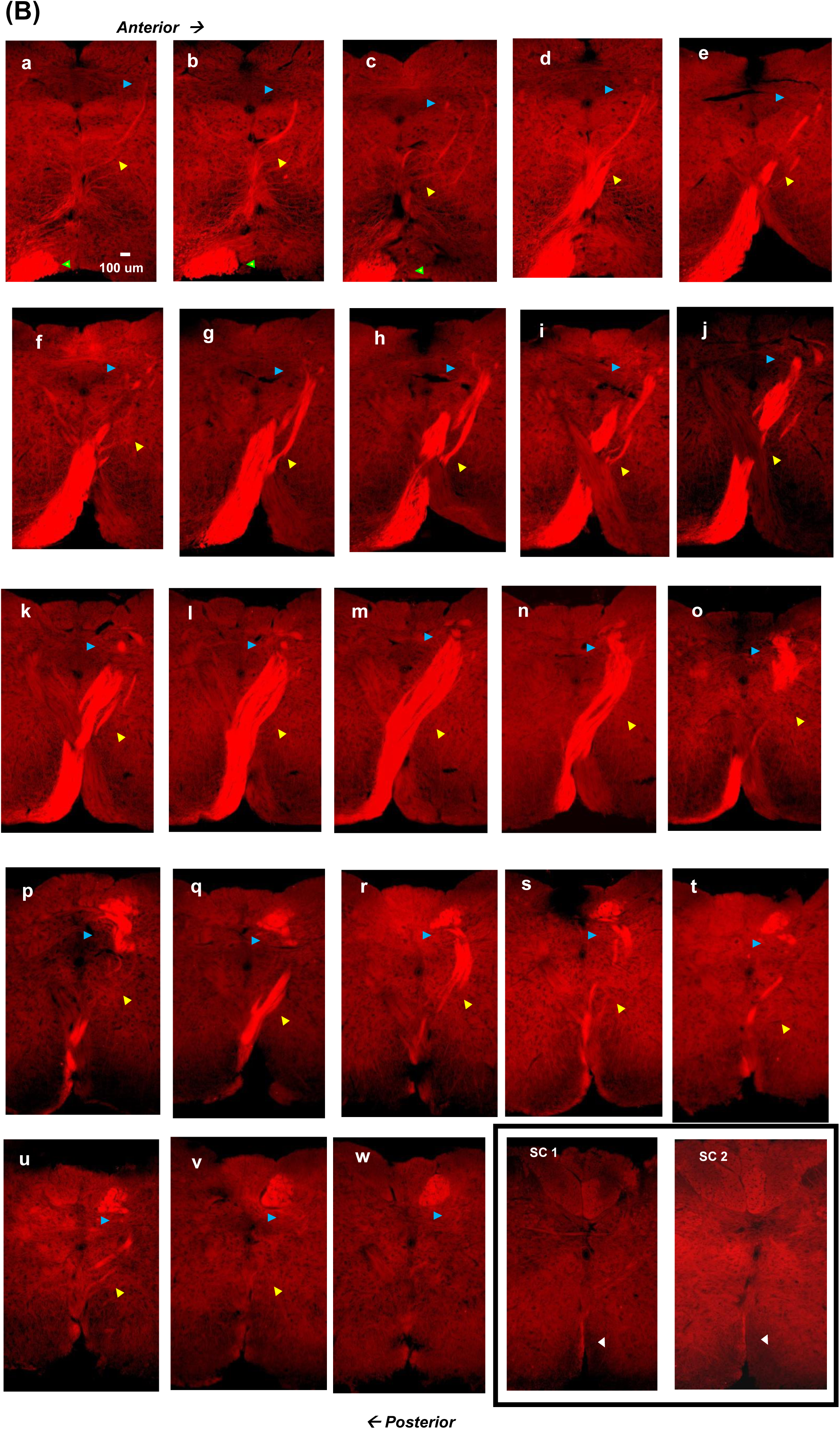
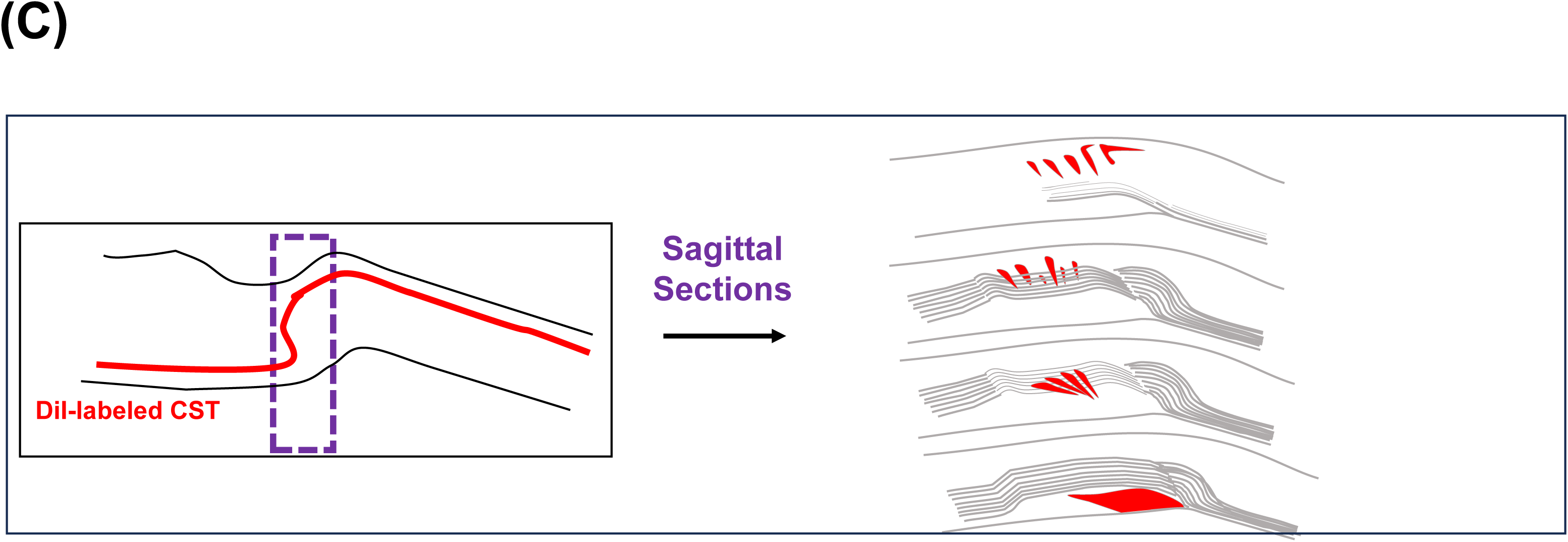
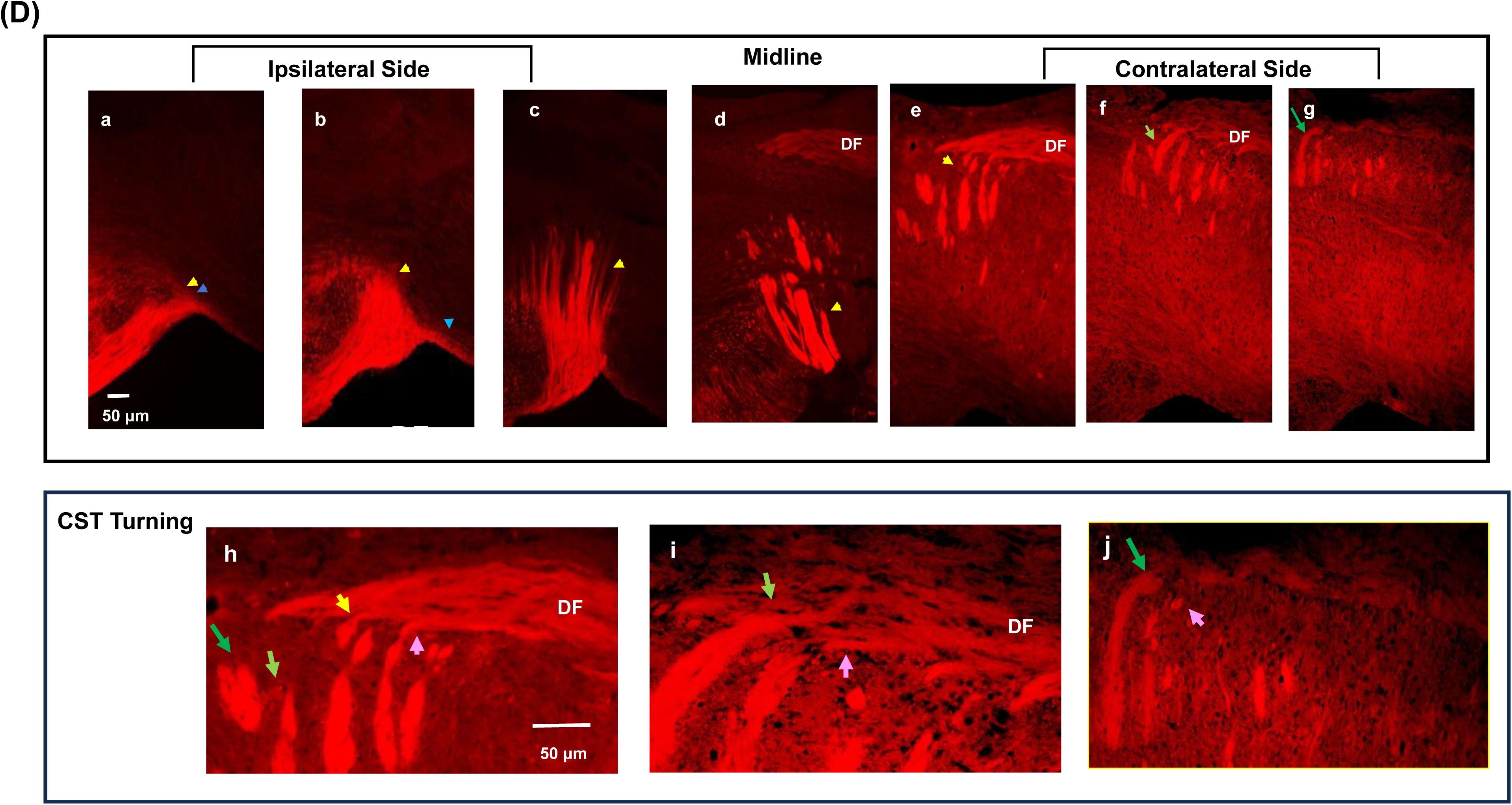
Coronal and Sagittal Analyses of CST decussation by DiI tracing. **(A)** A diagram that outlines the coronal analysis below. **(B)** CST decussation in 23 coronal sections (15 μm thick/each) by DiI tracing (n=10). In the most anterior sections, CST decussation is shown in faint DiI above the pyramidal tract (green arrows in a-c). In more posterior sections, CST (strong DiI) is a single tract ipsilaterally, but the fascicles separate in the midline and on the contralateral side (yellow arrows in d-n). The amount of CST fibers in the dorsal funiculus are shown in blue arrows. The uncrossing CST is in the medio-ventral side of spinal cord (SC 1-2). **(C)** A diagram that outlines the sagittal analysis. **(D)** CST decussation in 5 sagittal sections (15 μm thick/each) by DiI tracing (n=6). The crossing and uncrossing CSTs (yellow and blue arrows) travel together (a) but separate when the crossing CST moves antero-dorso-medially (b) and splits into 4 fascicle groups at midline (c, d). These fascicle groups further separate and turn to the dorsal funiculus (DF) in 3 contralateral sections (e-g). The turning areas are magnified (h-j). In the most medial section (h), the posterior fascicles turn to the ventral positions in DF (yellow arrows). In the next medial section (i), the more anterior fascicles turn to the more dorsal positions in DF (light green arrow). In the farthest medial section (j), the most anterior fascicles turn to the most dorsal positions in DF (dark green arrow). Within each section, the fascicles in the front turn more dorsally than the fascicles in the back and vice versa (green and pink arrows).

We next examined the distribution of non-crossing CSTs. To compensate for the low fascicle number, a higher amount of DiI solution was injected into the lateral side of one pyramidal tract in the neuraxis. As expected, the non-crossing CST is localized in the ventral/medial side of the spinal cord (white arrows in SC1- 2).

### 3. Sagittal Analysis

*Method (I): PKC Immunostaining:* N/A due to the lack of specificity. Method (II): Non-Invasive Anterograde Tracing

To assist in reading the sagittal analysis, we drew a diagram in Fig. 3C that outlines the experiments. Here, the DiI-labeled neuraxis was sectioned sagittally and analyzed without PKC-γ immunostaining. Figure 3D shows the results from ipsilateral to contralateral sections. In (a), the decussating CST (yellow arrow) and the non-crossing CST (blue arrow) travel together at the pyramidal tract. However, these two types of tracts separate at the junction between medulla and spinal cord where the decussating CST turns and moves antero-dorso-medially for decussation (b, c). During decussation at the midline, CST splits into fascicle groups (d). After decussation, these fascicles further separate at different locations while moving on the contralateral side (e-g). This final separation determines the turning position of the CS axons in the dorsal funiculus. This turning strategy can be clearly seen in the enlarged images of the dorsal funiculus (h-j). In the most medial section (h), only the posterior CS fibers turn and enter the ventral positions in the dorsal funiculus (yellow and the pink arrows in h). In contrast, the anterior CS fibers do not turn here (light and dark green arrows). In the next medial section (i), the next anterior CS fibers turn and enter the more dorsal positions in the dorsal funiculus (light green arrow). In the farthest medial section (j), the most anterior CS fibers turn and enter the most dorsal positions in the dorsal funiculus (dark green arrow). Beyond this section, there is no detectable CS fibers. This rule also applies to the fibers within each section in which the CS fibers in the front turn more dorsally than the ones behind (yellow/green vs pink arrows). Together, the sagittal analysis shows the utility of the oval structure in the separation of CS fibers. In conjunction with the turning strategy, the separated CS fibers can turn/occupy the correct position in the dorsal funiculus for proper limb control.

### The Major Components Associated with the Oval Structure

#### 1. Interior

To characterize the components in the oval structure, we first examined GFAP^+^ glial cells by co-immunostaining the horizontal sections for GFAP and PKC-γ (top diagram, Fig. 4A). We found two populations of GFAP^+^ cells (c). One population scattered in the oval structure (yellow arrows) whereas the second population expressed high amounts of GFAP and distributed at the leading edge of CSTs prior to entering the oval structure (white arrows in b, c). However, these cells gradually diminished as the numbers of CSTs in the oval structure increased (f, i, l, o). After decussation, the cells with weaker GFAP signals appeared at midline (pink arrow in q, r). Judging from this dynamic change, it was likely that the second population of GFAP^+^ cells was related to meninges.

**Figure 4:**
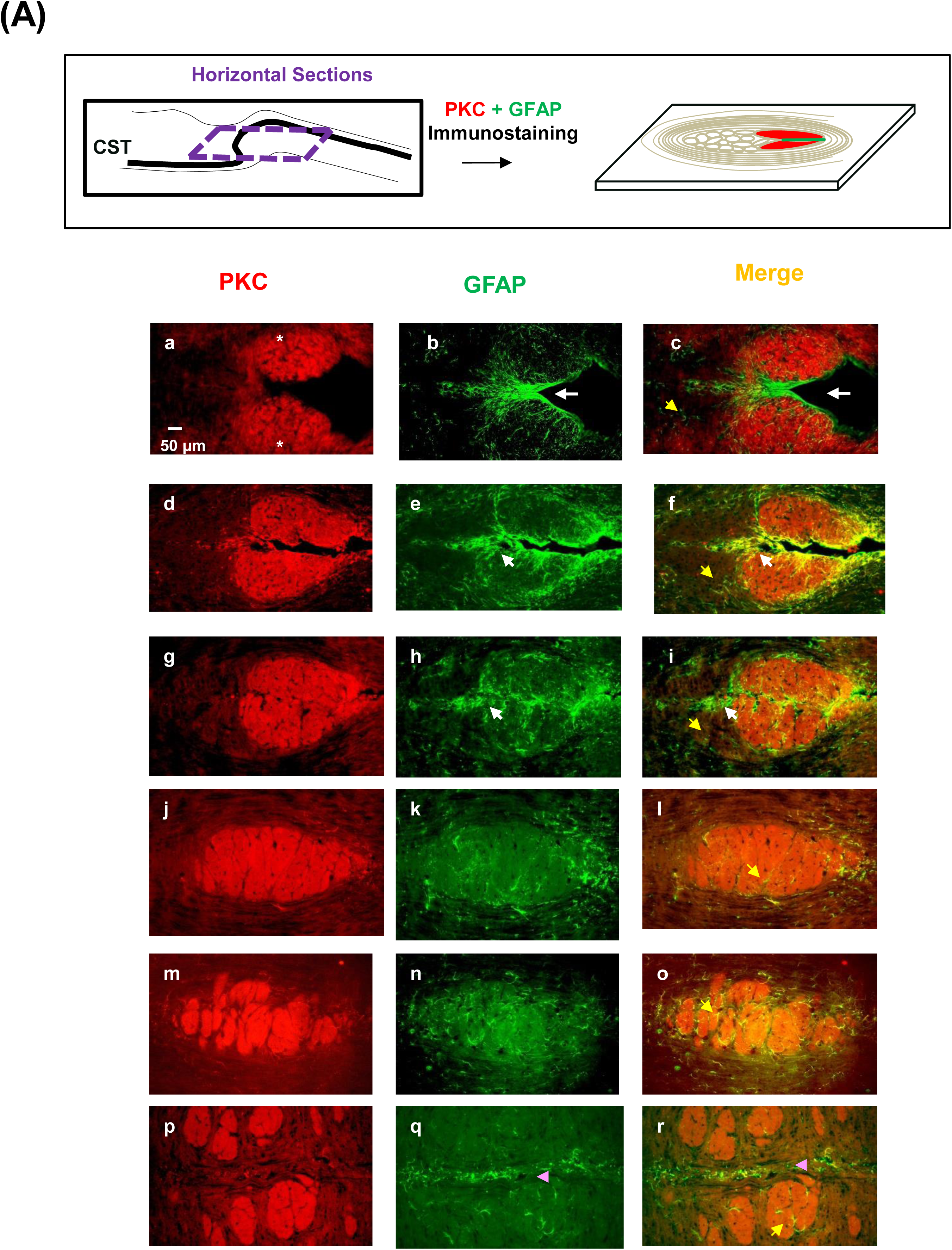
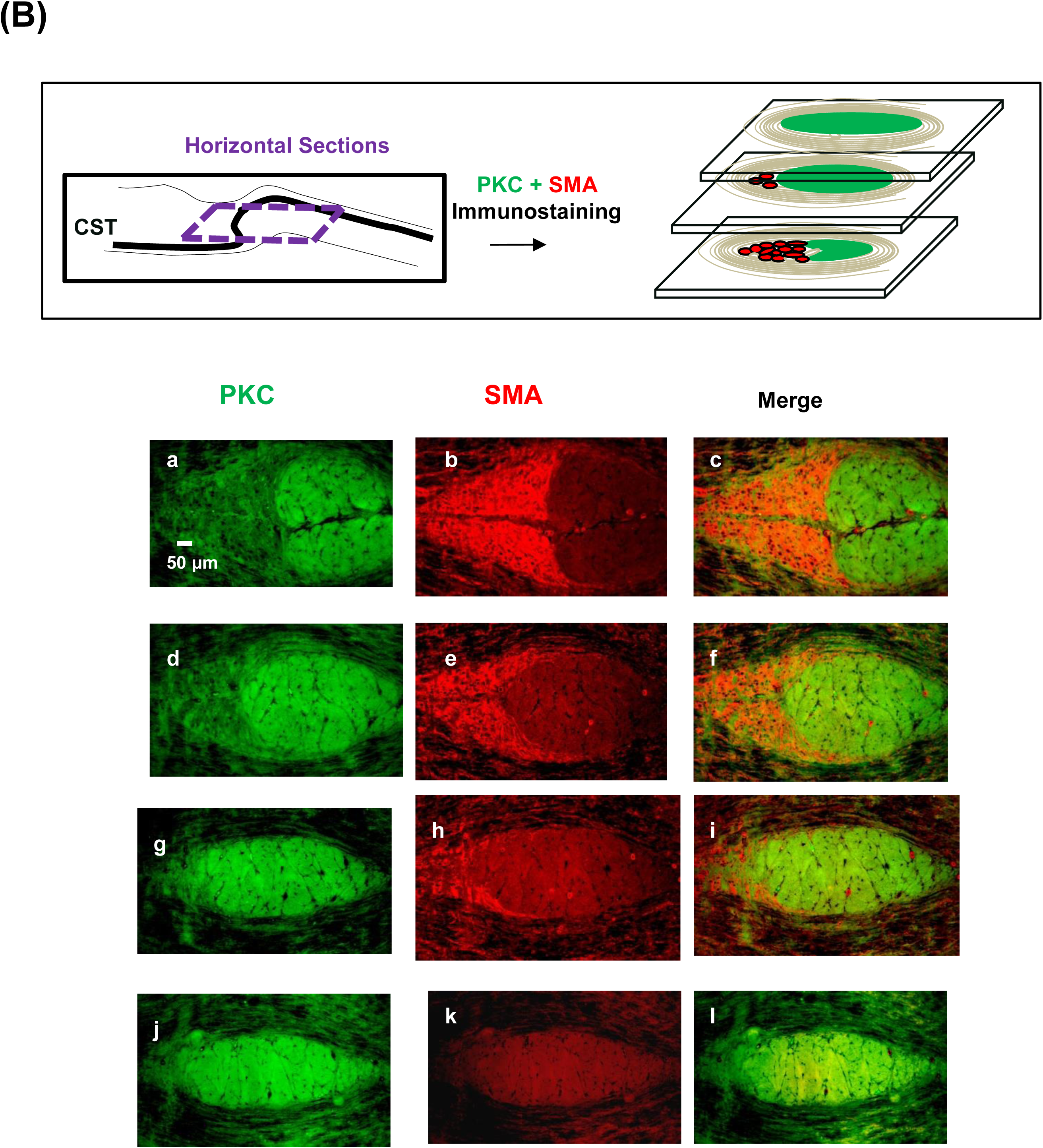
The major components in the oval structure. **(A)** Two populations of GFAP^+^ cells are in the oval structure when co-immunostained for GFAP and PKC-γ (n>15). One scatters in the oval structure (yellow arrows, c) and the other one is concentrated at the leading edge of CSTs (white arrows, b, c). The latter population gradually disappears during CST decussation (e, f, h, i, k, l). After decussation, the cells with weaker GFAP signals appear at midline (q, r). **(B)** The SMA^+^ cells in the oval structure (n>10) (top diagram). Co-immunostaining for SMA and PKC-γ shows the SMA^+^ cells in the oval structure gradually retreat to yield space for CSTs during decussation (c, f, i, l).

We then examined different isoforms of actin in the oval structure by co-immunostaining for different actin isoforms and PKC-γ (top diagram in Fig. 4B). Although β-actin was expressed in the oval structure, we chose smooth muscle actin (SMA) as the marker for the residential cells in this structure due to its higher specificity (supplementary Fig. S2). Figure 4B shows the SMA^+^ cells inside the oval structure gradually retreat and finally disappear to yield the space to accommodate CSTs for decussation (c, f, i, l).

These components (CSTs, GFAP^+^ cells and SMA^+^ cells) and their relative positions in the oval structure are shown by triple labelling in supplementary Fig. S3.

#### 2. Exterior

In addition to the interior components, we also characterized the cells that formed concentric layers surrounding the oval structure. We speculated these cells were non-CST nerve fibers. We first co-immunostained the horizontal sections for myelin basic protein (MBP) (green) and PKC-γ (red) (top diagram in Fig. 5A). Figure 5A shows the data from ventral to dorsal sections. The cells in the concentric layers express MBP. Moreover, the MBP^+^ concentric layers and the oval structure are present even before CSTs arrive (a-c) and enclose CSTs more thoroughly during decussation (f-l). However, after decussation, the oval structure and the MBP^+^ layers disappear (o, r).

**Figure 5:**
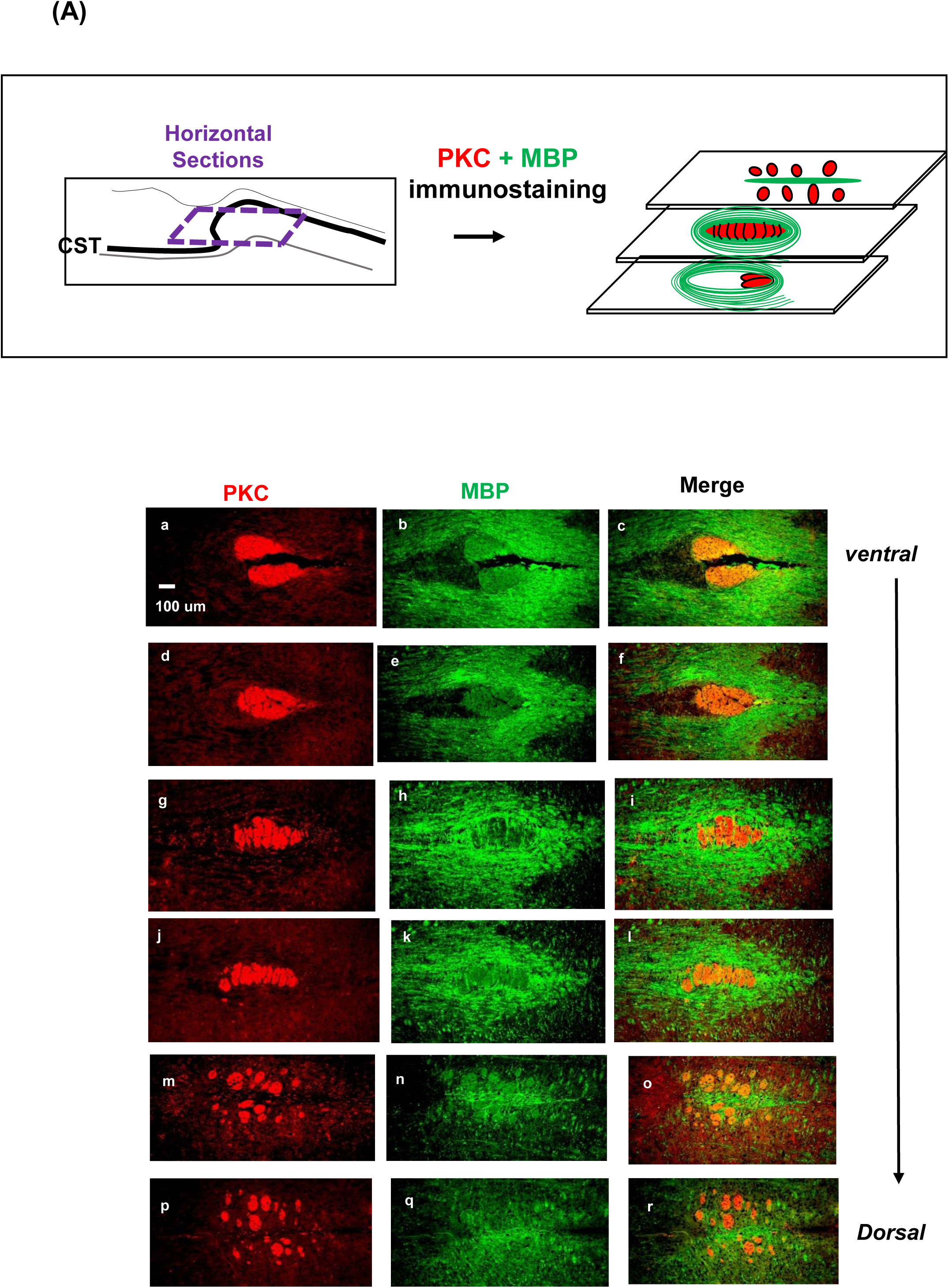
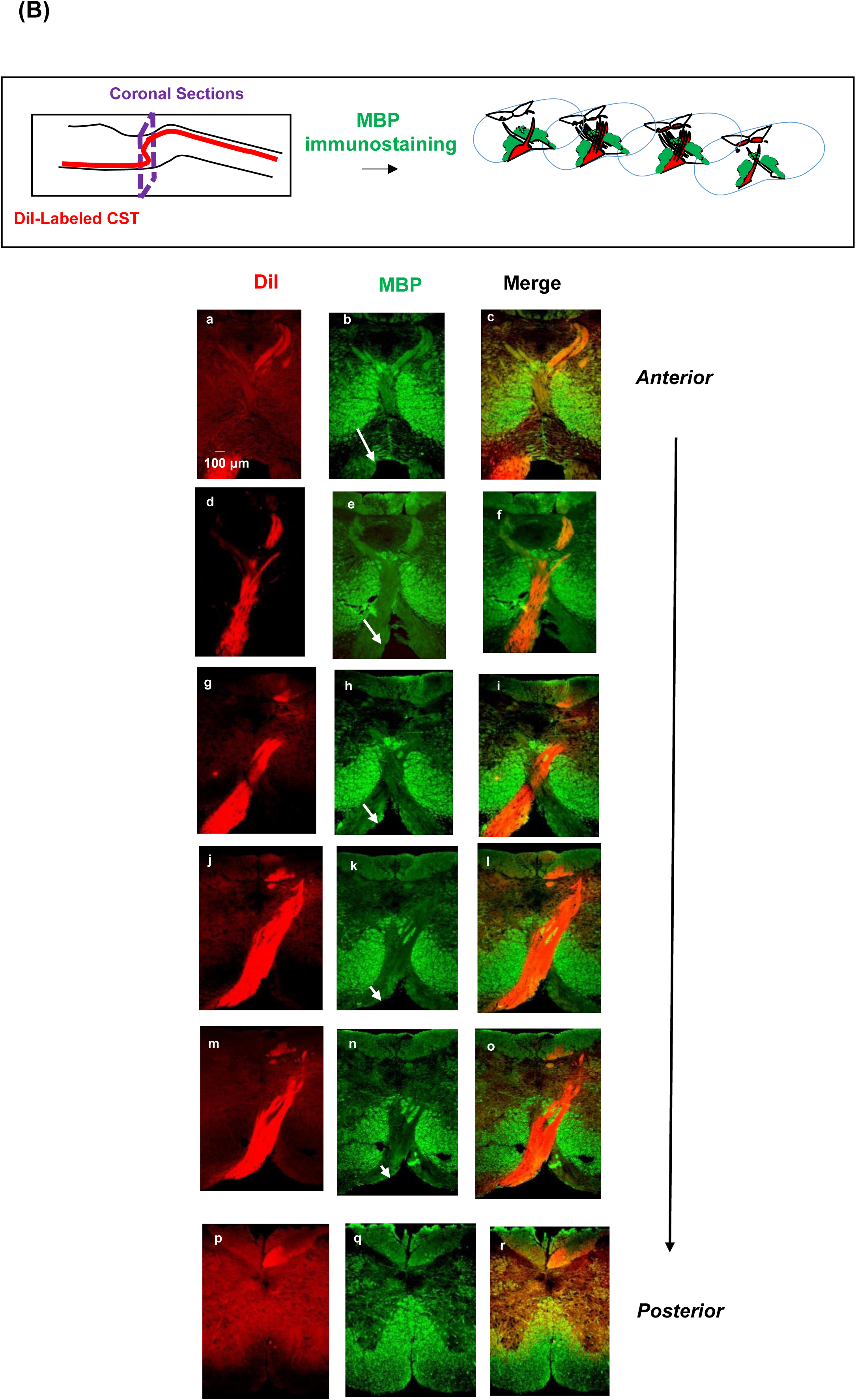
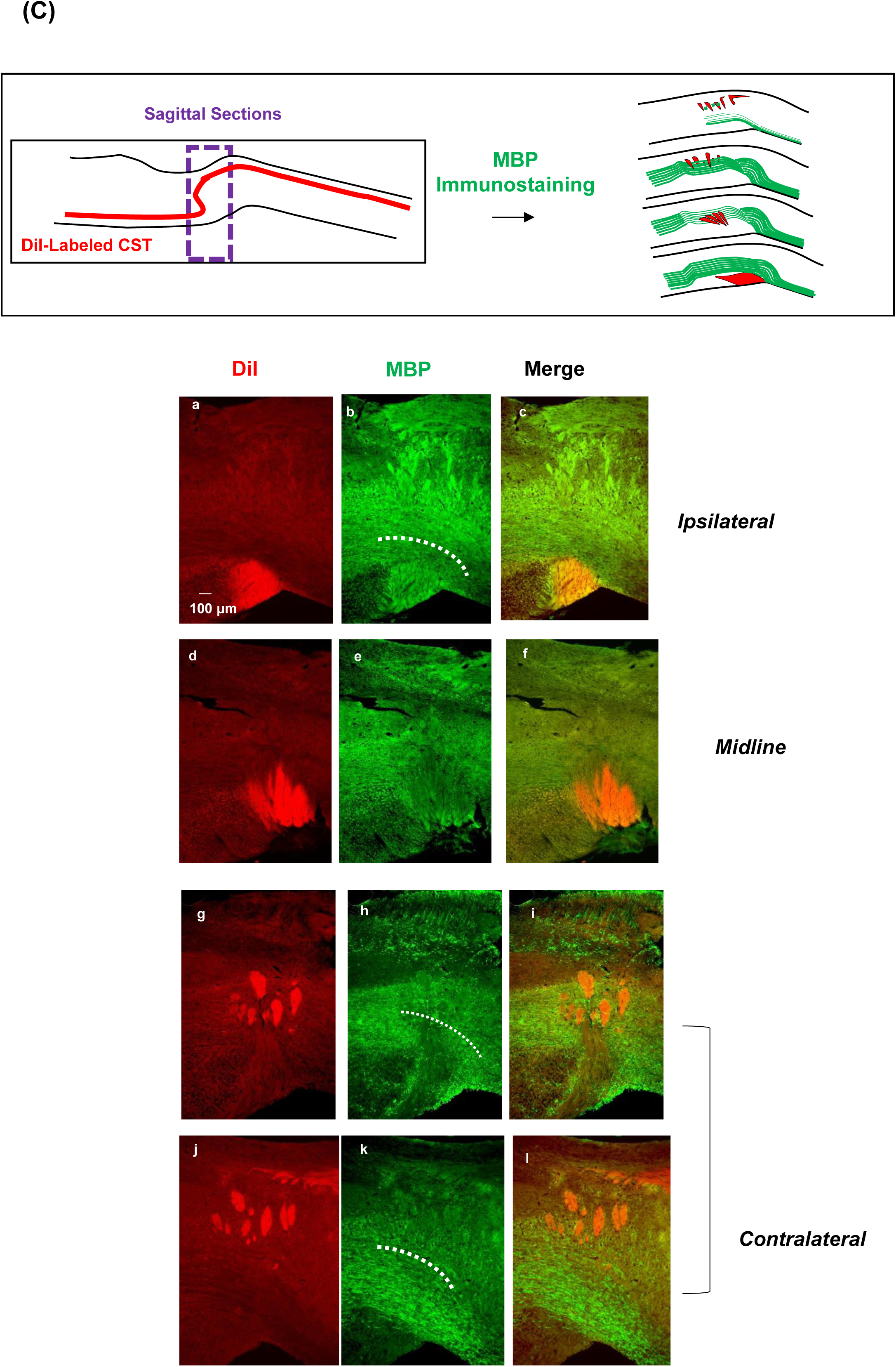
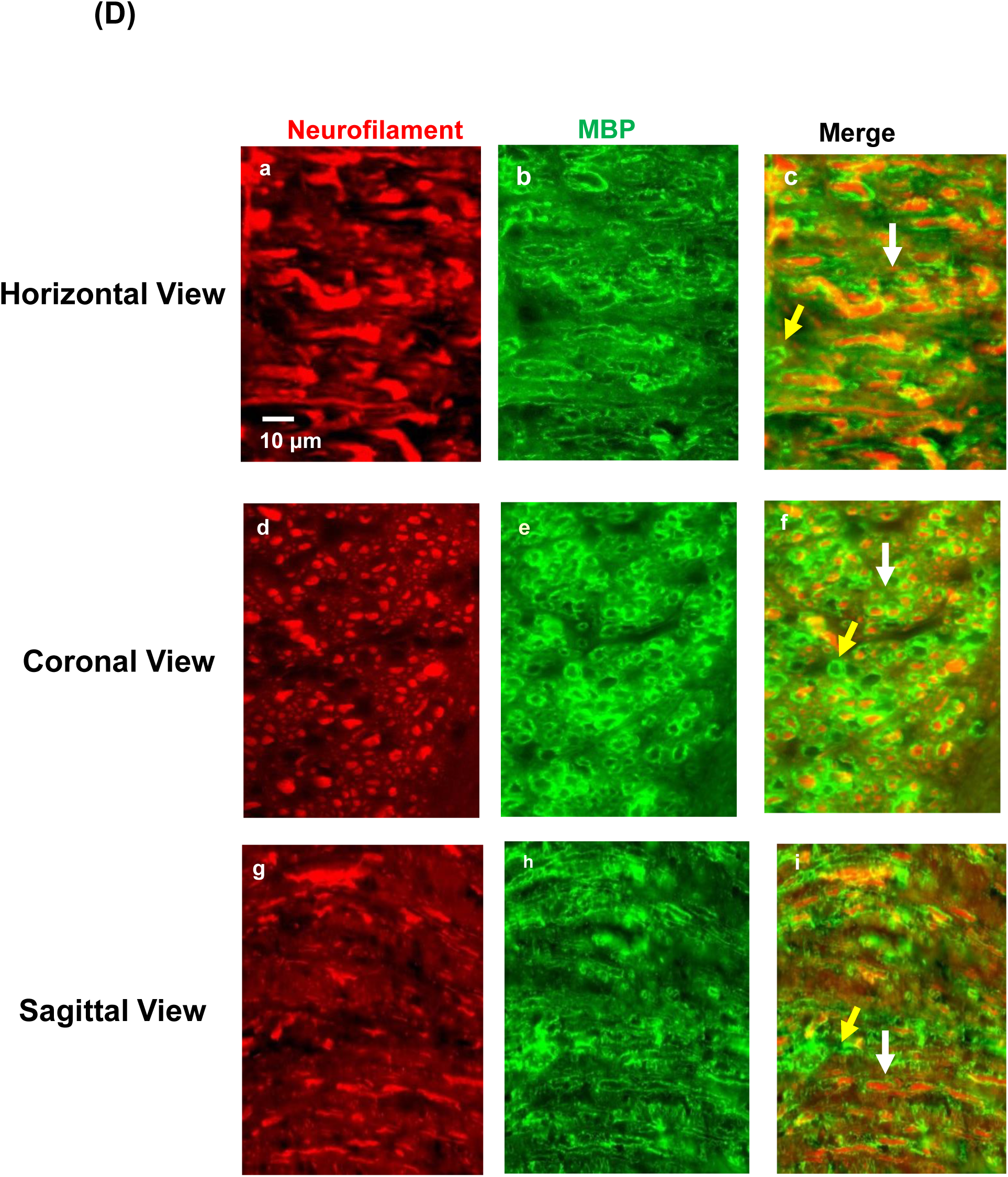
The major component surrounding the oval structure. The oval structure is surrounded by concentric layers of cells. Co-immunostaining for MBP (green) and PKC-γ (red) show these cells express MBP in 3 anatomical planes (top diagrams). **(A)** Horizontal analysis: The MBP^+^ cells form concentric layers that surround the oval structure before CSTs arrive (a-c) and enclose the oval structure more thoroughly during CST decussation (f.i,l). After decussation, the MBP^+^ cells and the oval structure disappear (o.r). **(B)** Coronal analysis: The MBP^+^ cells sandwich CSTs at midline i.e., during decussation (c,f,i,l,k) and change their positions to accompany CSTs, from more dorsal/lateral positions down to the ventral/medial positions (white arrows, b,e,h,k,n). **(C)** Sagittal analysis: The MBP^+^ cells appear as a green belt in medulla but gradually descend to the ventral side near spinal cord (white dashed line in the MBP column). This was not obvious in the midline (f, i). **(D)** Co-immunostaining for MBP and neurofilament shows each MBP^+^ cell circles a neurofilament under high magnification (c.f.i), suggesting these cells are non-CST nerve fibers.

The appearance of these MBP^+^ cells was also examined coronally and sagittally (Fig. 5B and C). The DiI-labeled neuraxis was sectioned coronally or sagittally and immunostained for MBP (top diagram in Fig. 5B and C). Figure 5B summarizes the coronal data from anterior to posterior sections. In these sections, the MBP^+^ cells sandwich the CSTs at midline (green in c,f,i,l,o).

However, the position of the MBP^+^ cells change during CST decussation. At the beginning of decussation, the MBP^+^ cells are above the ventral border and lateral to the midline (white arrow in b). However, as decussation progresses, the MBP^+^ cells gradually descend diagonally to the ventro-medial positions that is indicated by changes in the length of white arrows in panels e, h, k and n. Figure 5C summarizes the sagittal data from ipsilateral to contralateral sections. Before decussation in an ipsilateral section (dashed white curve in b), the MBP^+^ cells appear like a green belt in medulla but tilt down in the spinal cord. However, during decussation in the midline sections, this green belt cannot be seen clearly (e,f). After decussation in the contralateral sections, this green belt re-appeared in medulla and tilted down to the ventral side near spinal cord (dashed white curve in h, k, i, l). These results suggested the MBP^+^ cells which surrounded the oval structure in medulla merged with the white matter on the ventral side of spinal cord.

To confirm that these MBP^+^ cells were non-CST neural fibers, we analyzed the expression of MBP and neurofilaments in 3 anatomical planes. Figure 5D shows the results under high magnification, in which each MBP^+^ cell circles around a neurofilament (white arrows in a-c, d-f, g-i). However, due to imaging limitations, the signals from the neurofilament are weak in a small number of MBP^+^ cells because these cells were in deeper layers (yellow arrows). We also confirmed these results using different antibodies against MBP and tubulin 3b.

## Discussion

In this report, we identified a new pathway for CST decussation. This pathway is illustrated from the views of 3 anatomical planes in Fig. 6 (top). In contrast to the current belief, we found CSTs did not decussate at the junction between medulla and spinal cord. Instead, CSTs decussated in an oval structure anterior to this junction. In this oval structure, each CST split into 4 fascicle groups. Each group then interdigitated with the corresponding groups from the opposite CST to cross the midline. These data suggested CSTs did not decussate as a single tract, but as fascicle groups. The significance of this pathway was revealed by the consequences after decussation. Once on the contralateral side, the 4 fascicle groups changed direction, moving toward spinal cord. While moving, the CS fibers in each fascicle group separated at different locations. This separation determined the turning position of each CS fiber in the dorsal funiculus (bottom, Fig. 6). The CS fibers that separated anteriorly (A/P coordinates) turned laterally (M/L coordinates) and occupied the dorsal positions (D/V coordinates) in the dorsal funiculus and vice versa. This turning process is essential for proper limb control. Together, our new pathway suggests CST decussation is not merely crossing the midline. The use of the oval structure allows CS fibers to cross the midline and separated at different locations and subsequently turned into the correct positions in the dorsal funiculus for proper limb control.

**Figure 6:**
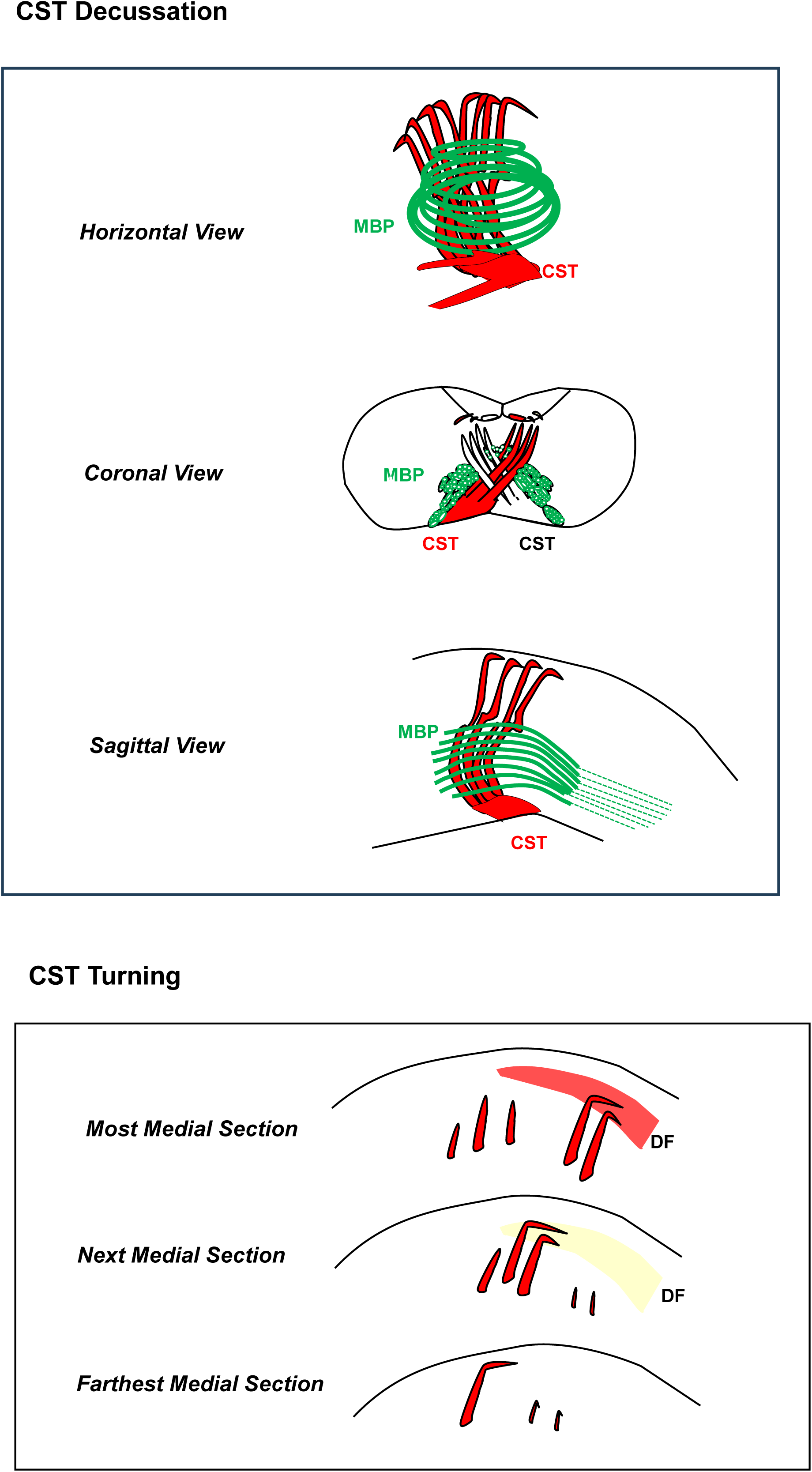
A new model for CST decussation. The new model for CST decussation is illustrated from 3 anatomical planes **(top)** which show the different views of CSTs, the oval structure, the surrounding MBP^+^ nerve fibers and the resulting separation of CST fascicles. The separated CST fascicles followed a turning strategy to turn and enter the dorsal funiculus (DF). This turning strategy is illustrated in a sagittal view **(bottom).** Between sections, there was a high correlation between the A/P coordinates of a fascicle, the M/L section to turn and the D/V turning positions in the dorsal funiculus. This rule also applied to the fascicles within the same section.

In fact, the separation of CS fibers after decussation has been indirectly observed before. In previous reports, this separation appeared as the co- existence of multiple CST fascicles but this phenomenon was overlooked ^4–6^.

Moreover, our mapping data on M1 supported this new pathway. Our preliminary data suggested the CST fibers from the hindlimb areas in M1 were the first fibers to enter the oval structure for decussation and therefore became the anterior fascicles after decussation. These fibers then turned and entered the dorsal positions in the dorsal funiculus. The same logic also applied to the CST fibers from the forelimb areas in M1. These mapping experiments are currently in progress.

## Materials and Methods

The animal experiments in this study were approved by IACUC at SUNY Downstate Medical Center. All of the animal experiments were performed in accordance to the guidelines and regulations of this approved IACUC protocol. Moreover, all data on animal research are reported in accordance to the ARRIVE guidelines.

### Stereotaxic DiI Tracing *in vitro*

DiI crystals were prepared as a solution (1% w/v). Please note, the commercially available DiI solutions are too dilute and therefore not suitable for this method. To prevent clogging, a filamented borosilicate glass capillary was used for injection (A-M system, WA). The DiI solution was backfilled into the microfilament in a borosilicate glass capillary and the tip of this glass was manually broken for injection. The neuraxis above the mid-segments of the spinal cord was isolated from adult mice (6-8 wks old C57BL/6 from Charles River, MA) postmortem. Prior to isolating the neuraxis, an 18G needle penetrated the Bregma point to mark this position on the brain. The isolated neuraxis was fixed with 4% paraformaldehyde. During DiI injection, the fixed neuraxis was placed in a 35 mm culture dish which was half filled with PBS+0.02% sodium azide (Becton Dickinson, NJ) and the glass capillary was mounted to a stereotaxic apparatus. The center point of the fissure between olfactory bulb and the brain of the neuraxis is counted as 0. DiI was injected into one of the pyramidal tracts (below pons AP: 0.2-0.5, ML: 0.4-0.6, DV: 0.2-0.4) or M1 (AP: 1-2.2, ML:0.7-1.4, DV: 1.0-1.4) of the neuraxis unilaterally *in vitro*. After injection, the neuraxis was kept at room temperature for 2-3 months before analysis.

### Immunostaining

The brain section was incubated with the primary antibody at 4^0^C overnight, followed by PBS washing and then incubated with FITC-labeled or Cy3- labeled secondary antibody. The antibodies used were PKC-γ (Dr. Sactor’s lab/ Genescript), SMA (Abcam), GFAP (Aves Lab), MBP (Aves Lab/Sigma) and Neurofilament (Dako).

## Acknowledgements

We thank Drs. Chang-Chi Hsieh, Ilham Muslimov, Andrew Tcherepanov, Elena Nikulina and Jenny Libien for the reagents. We thank Dr. Joshua Sanes and Ms. Seto Chice for reviewing the early drafts of this manuscript.

## Author Contributions

Conceptualization: HLL

Methodology: HLL

Investigation: HLL

Visualization: HLL

Funding acquisition: HLL, NL, MA

Project administration: HLL

Writing – original draft: HLL

## Data and Material Availability

All data are available in the main text or the supplementary materials.

## References

1 Welniarz, Q., Dusart, I. & Roze, E. The corticospinal tract: Evolution, development, and human disorders. Dev Neurobiol 77, 810–829 (2017). 10.1002/dneu.22455

2 Iordanova, R. & Reddivari, A. K. R. in StatPearls (2024).

3 Dottori, M. et al. EphA4 (Sek1) receptor tyrosine kinase is required for the development of the corticospinal tract. Proc Natl Acad Sci U S A 95, 13248–13253 (1998). 10.1073/pnas.95.22.13248

4 Welniarz, Q. et al. Non cell-autonomous role of DCC in the guidance of the corticospinal tract at the midline. Sci Rep 7, 410 (2017). 10.1038/s41598-017-00514-z

5 Aizawa, S. et al. Abnormal Pyramidal Decussation and Bilateral Projection of the Corticospinal Tract Axons in Mice Lacking the Heparan Sulfate Endosulfatases, Sulf1 and Sulf2. Front Mol Neurosci 12, 333 (2019). 10.3389/fnmol.2019.00333

6 Faulkner, R. L. et al. Dorsal turning of motor corticospinal axons at the pyramidal decussation requires plexin signaling. Neural Dev 3, 21 (2008). 10.1186/1749-8104-3-21

