## Supplementary Figures for "A New Pathway for the Decussation of Corticospinal Tracts"

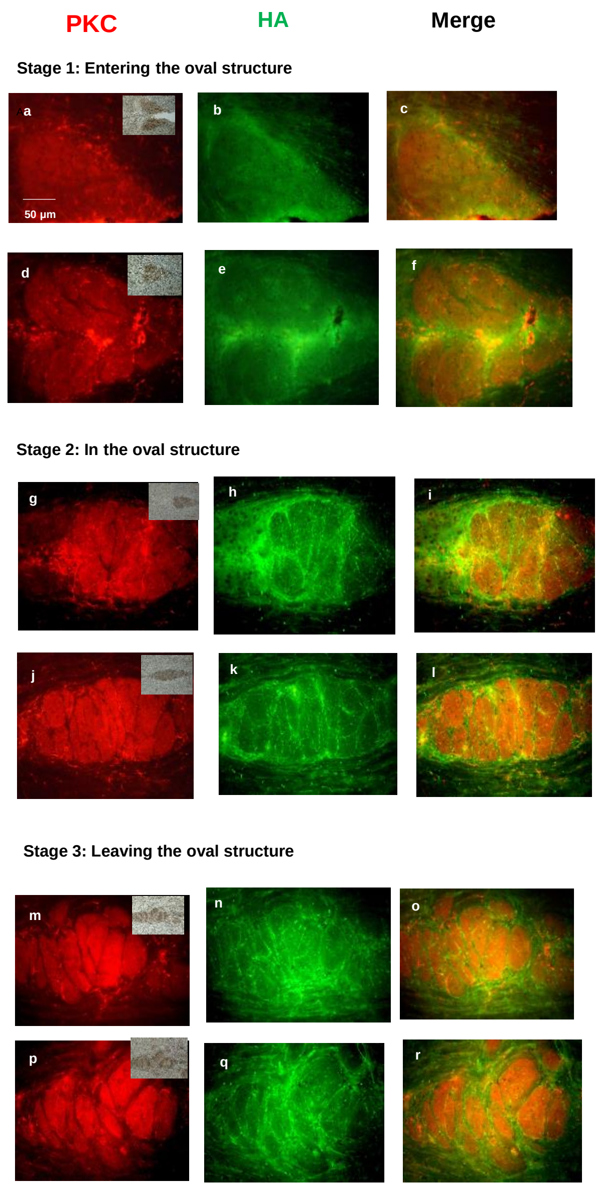


Supplementary Figure S1

**Supplementary Figure S1: Increased extracellular space in the oval structure**

The horizontal sections were co-immunostained for hyaluronic acid (HA) and PKC-γ. The results are presented from ventral to dorsal sections. All sections show both CSTs except the most ventral section in which only one CST is shown (a-c). The expression of HA is low before decussation (c,f, stage 1), but increases when both CSTs are in the oval structure (i, l, stage 2). This increased expression of HA persists after decussation (o,r, stage 3).

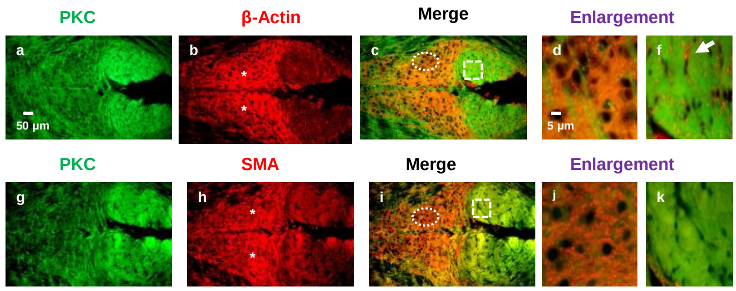


Supplementary Figure S2

**Supplementary Figure S2: Punctate expression of β-actin and SMA**
The residential cells in the oval structure express both β-actin and SMA (b,c,h,i). Both β-actin and SMA show punctate staining pattern in the magnified images (d,j for the circled areas in c,i). However, when magnifying the area of CSTs (f, k for the square area in c,i), β-actin, but not SMA is found in the intercellular space (arrow in f). This suggests that β-actin is not a specific maker for the oval structure.

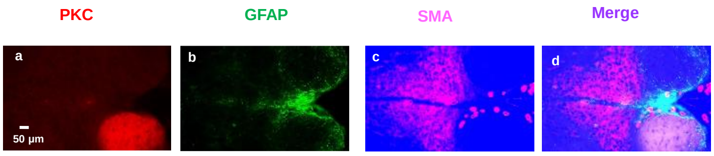


Supplementary Figure S3

**Supplementary Figure S3: The relative positions of CST, GFAP$^+$ cells and SMA in the oval structure**

The CST was labeled by DiI injection (red, a). The horizontal sections of the DiI-labeled neuraxis were immunostained for GFAP (green, b) and SMA (bright pink, c). Upon merge (d), CST becomes light pink, GFAP is light blue and SMA is bright pink.
